# Calcineurin-mediated regulation of growth-associated protein 43 is essential for neurite and synapse formation and protects against α-synuclein-induced degeneration

**DOI:** 10.1101/2025.01.13.632848

**Authors:** Sofia Zaichick, Ekaterina Grebenik, Gabriela Caraveo

## Abstract

**Introduction:** Elevated calcium (Ca^2+^) levels and hyperactivation of the Ca^2+^-dependent phosphatase calcineurin are key factors in α-synuclein (α-syn) pathobiology in Dementia with Lewy Bodies and Parkinson’s Disease (PD). Calcineurin activity can be inhibited by FK506, an FDA-approved compound. Our previous work demonstrated that sub-saturating doses of FK506 provide neuroprotection against α-syn pathology in a rat model of α-syn neurodegeneration, an effect associated with the phosphorylation of growth-associated protein 43 (GAP-43).

**Methods:** To investigate the role of GAP-43 phosphorylation, we generated phosphomutants at the calcineurin-sensitive sites and expressed them in PC12 cells and primary rat cortical neuronal cultures to assess their effects on neurite morphology and synapse formation. Additionally, we performed immunoprecipitation mass spectrometry in HeLa cells to identify binding partners of these phosphorylation sites. Finally, we evaluated the ability of these phosphomutants to modulate α-syn toxicity.

**Results:** In this study, we demonstrate that calcineurin-regulated phosphorylation at S86 and T172 of GAP-43 is a crucial determinant of neurite branching and synapse formation. A phosphomimetic GAP-43 mutant at these sites enhances both processes and provides protection against α-syn-induced neurodegeneration. Conversely, the phosphoablative mutant prevents neurite branching and synapse formation while exhibiting increased interactions with ribosomal proteins.

**Discussion:** Our findings reveal a novel mechanism by which GAP-43 activity is regulated through phosphorylation at calcineurin-sensitive sites. These findings suggest that FK506’s neuroprotective effects may be partially mediated through GAP-43 phosphorylation, providing a potential target for therapeutic intervention in synucleinopathies.

## Introduction

Misfolding of the small lipid binding protein α-synuclein (α-syn), plays a central role in a group of neurological diseases collectively known as synucleinopathies (Alafuzoff and Hartikainen, 2017). These encompass Parkinson’s Disease (PD), Multiple Systems Atrophy and two prominent Lewy Body Dementias: Dementia with Lewy Bodies (DLB) and Parkinson’s Disease Dementia (PDD). Using a diverse array of model systems, we and others have found that α-syn leads to a pathological increase in cytosolic Ca^2+^ and a subsequent hyperactivation of calcineurin (CaN), the highly evolutionary conserved Ca^2+^-Calmodulin-dependent serine/threonine phosphatase resulting in cell death (Chan et al., 2007; Dufty et al., 2007; Surmeier et al., 2017; Yuan et al., 2013; Caraveo et al., 2017; Guzman et al., 2010; Hurley et al., 2013; Surmeier et al., 2010; Goldberg et al., 2012; Surmeier et al., 2016; Caraveo et al., 2014; Martin et al., 2012; Burbulla et al., 2017). Importantly, we found that reducing CaN activity—either genetically or with sub-saturating doses of the CaN-specific inhibitor FK506 (also known as Tacrolimus)—rescues cells from the toxic effects of α-syn *in vitro* and in a large preclinical model (Caraveo et al., 2014; Caraveo et al., 2017).

Growth-associated protein-43 (GAP-43) is a nervous system-specific protein (Karns et al., 1987) that plays a crucial role in neuronal development, particularly in growth cone pathfinding (Frey et al., 2000). Beyond development, GAP-43 remains essential in adulthood, facilitating neuronal regeneration after injury and contributing to synaptic plasticity (Holahan, 2017). Its ability to promote growth cone formation is attributed to its regulation of both actin dynamics and vesicle recycling at presynaptic terminals (Denny, 2006; Korshunova et al., 2008). The activity of GAP-43 can be regulated by multiple mechanisms, including transcriptional expression, membrane association via palmitoylation, and phosphorylation, modulated by Ca^2+^ signaling—specifically, through phosphorylation by Protein Kinase C (PKC) and dephosphorylation by CaN at the S41 site (Chan et al., 1986; Skene, 1989; Liu and Storm, 1989; Nielander et al., 1990). Emerging evidence links GAP-43 to PD. Lower GAP-43 levels have been detected in cerebrospinal fluid (Sjogren et al., 2000) and postmortem substantia nigra pars compacta tissue from PD patients (Saal et al., 2017). Consistent with these findings, an *in vitro* scratch lesion model demonstrated that α-syn mutants associated with early-onset PD failed to regenerate axons, a defect which correlated with lower GAP-43 expression (Tonges et al., 2014).

In our study, we discovered a novel function for the S86 and T172 phosphorylation sites on GAP-43, which are regulated by CaN under α-syn conditions. We hypothesized that FK506-mediated phosphorylation of GAP-43 at S86 and T172 enhances neurite branching and synapse formation, thus providing neuroprotection against α-synuclein toxicity. Using phosphomutants in cell lines and primary cortical neurons, we demonstrated that phosphorylation at these sites is essential for both neurite branching and synapse formation. Specifically, the phosphomimetic mutant of GAP-43 (S86E,T172E), which represents GAP-43 status under α-syn neuroprotective conditions due to partial inhibition of CaN, promotes neurite outgrowth and mature synapses. The phosphoablative mutant of GAP-43 (S86A,T172A) on the other hand, which represents GAP-43 status under α-syn neurotoxic conditions due to constitutive CaN activity, does not promote neurite outgrowth and yields immature synapses. We found that the α-syn– and CaN-sensitive phosphorylation sites serve as a scaffold for proteins enriched in ribosomal function. Furthermore, overexpression of the GAP-43 phosphomimetic mutant (S86E,T172E), rescues α-syn–induced neuronal toxicity by increasing ATP levels, neuron viability, and PSD95 (postsynaptic density protein-95) expression. In contrast, the GAP-43 phosphoablative mutant (S86A,T172A), does not mitigate α-syn–mediated neurodegeneration. Collectively, these findings unveil a new mechanism by which CaN regulates GAP-43 activity. Furthermore, they provide a new mechanism by which FK506 exerts its neuroprotective effects, by modulating GAP-43 phosphorylation, thereby mitigating α-syn proteotoxicity.

## Results

### 1. GAP-43 phosphosites S86 and T172 are regulated by CaN and play a key role in neurite length and branching in PC12 cells

FK506 is a well-established drug and is already in widespread clinical use at high doses to suppress the rejection of organs in transplant patients, a process in which CaN also plays a critical role (Tron et al., 2018). Importantly, we found that sub-saturating doses of FK506 have excellent, long lasting brain penetrance and rapid clearance from the blood preventing potential adverse side effects (Caraveo et al., 2017). With a weekly dosing regimen, we embarked into a large *in vivo* study using a rat model of PD based on a unilateral injection of α-syn into Substantia Nigra Pars Compacta (SNc) (Caraveo et al., 2017). For the analysis, animals were stratified based on their final content of FK506 in the brain. We found that only sub-saturating doses of FK506 (<5ng/ml) were enough to prevent neurodegeneration by ameliorating the deficits in motor behavior, dopamine and dopamine transporter levels in the striatum caused by α-syn (Caraveo et al., 2017). To identify the substrates dephosphorylated by CaN associated with the toxic *vs* protective responses we took an isobaric unbiased phospho-proteomic approach using Tandem Mass Tagging (TMT) in combination with mass spectrometry (MS) from striatal tissue of these animals (Caraveo et al., 2017). We detected 51 of 3,526 phosphopeptides after correction for protein abundance, that changed significantly when comparing α-syn animals to the control. Among these, half of the phosphopeptides were significantly hypo-phosphorylated in α-syn animals compared to the controls (Figure 1A). Given that CaN is a phosphatase highly activated under elevated α-syn levels, we focused our analysis on the hypo-phosphorylated peptides. Four of these phosphopeptides restored their phosphorylation to control levels upon treatment with sub-saturating doses of Tacrolimus. These peptides belonged to only two proteins: GAP-43 (also known as F1, neuromodulin, B-50, G50, and pp46) and BASP1 (also known as NAP-22 and CAP-23) (Figure 1A,B).

**Figure 1.**
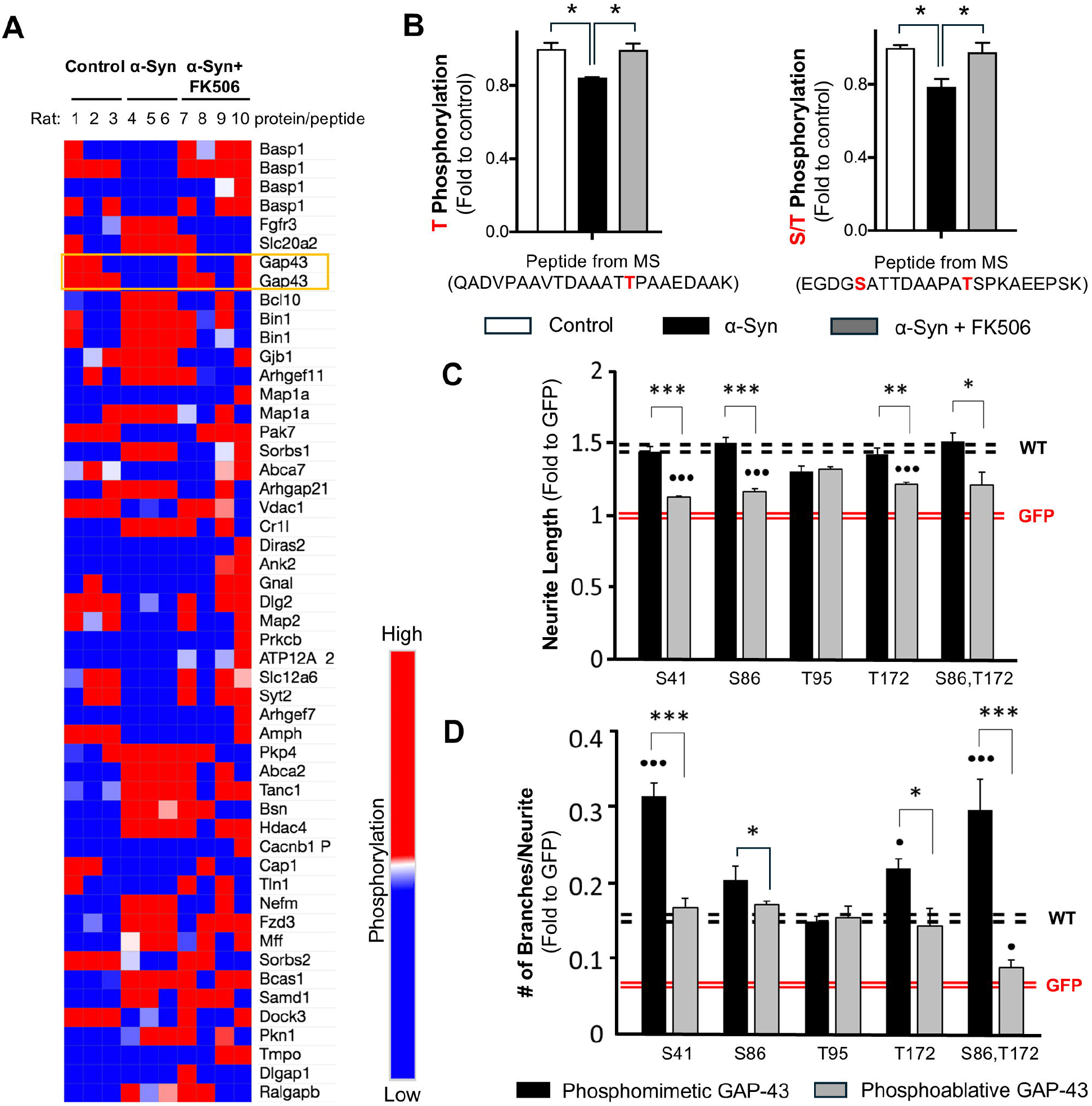
Calcineurin-dependent phosphosites S86 and T172 of GAP-43 contribute to neurite branching in PC12 cells. **(A)** A heat map representing phosphoproteomic hits identified in the rat striatum after correction for protein abundance, showing statistically significant differences between control and α-synuclein-expressing rats (Caraveo et al., 2017). Each row represents a protein from which the corresponding phosphorylated peptide was detected. Peptides highlighted in orange indicate GAP-43 phosphorylation sites that were restored to control levels in α-synuclein-expressing rats treated with sub-saturating doses of FK506 **(B)** Phosphopeptides from GAP-43 rescued with <5 ng/mL FK506 treatment in α-syn rat striatum. The identified phosphorylation sites are highlighted in red. Data represent n=3 rats. *p< 0.05 (two-tailed t-test). **(C-D)** PC12 cells were co-transfected with GFP and empty vector or GAP-43 in Wild type (WT) or the indicated phosphomutant form. The analysis includes measurements of the longest neurite **(C)** and the number of branches **(D)** from >300 cells across 4 independent experiments. All experiments are normalized relative to GFP control (red dashed lines; each line represents +/-SEM); WT GAP-43 serves as reference (black lines; each line represents +/-SEM). *p< 0.05 comparison between phosphomimetic and phosphoablative mutants and • p< 0.05 comparison between phosphomutants and WT; One-way Anova with post hoc Tukey’s test.

GAP-43 and BASP1 are functionally related proteins essential for neuronal development due to their roles in growth cone pathfinding (Frey et al., 2000). Moreover, these proteins play critical roles in neuronal regeneration and plasticity following injury (Holahan, 2017). To assess the functional significance of the phosphorylation sites identified through our unbiased TMT-MS approach (S86, T95, and T172 of GAP-43) in neurite growth, we generated phosphomimetic and phosphoablative mutants of these CaN-sensitive sites in GAP-43. We first tested the GAP-43 phosphomutants in PC12 cells, a rat pheochromocytoma cell line capable of differentiating into neuronal-like cells upon exposure to soluble Nerve Growth Factor or transfection with neuronal growth proteins such as GAP-43 (Greene and Tischler, 1976). PC12 cells were co-transfected with GFP (green fluorescent protein, to visualize neurites) and either wild-type (WT) GAP-43 or S86, T95, and T172 phosphomutant forms. As a positive control, we included the previously identified CaN-sensitive site S41. As a negative control we included GFP with empty vector. Cells were imaged 24 hours post-transfection and analyzed for neurite length and branching (Supplementary Figure 1). As expected, WT GAP-43 and the phosphomimetic-S41 mutant increased neurite length compared to GFP alone (Figure 1C), whereas the phosphoablative-S41 mutant had diminished effect. Among the three retrieved phosphosites, only the phosphomimetic-S86 and phosphomimetic-T172 mutants enhanced neurite length to levels comparable to WT GAP-43, whereas their phosphoablative counterparts had diminished effect (Figure 1C). These data suggest that phosphorylation at S41, S86 and T172 is required to promote neurite elongation.

We then tested the effect of these mutants in their ability to modulate neurite branching. As expected, the phosphomimetic-S41 mutant increased neurite branching compared to GFP alone and to WT GAP-43, whereas the phosphoablative-S41 mutant had diminished effect (Figure 1D). Similar to the neurite length results, only the phosphomimetic-S86 and phosphomimetic-T172 mutants enhanced neurite branching above WT GAP-43 and GFP alone, whereas their phosphoablative counterparts had diminished effect (Figure 1D). Notably, the double phosphomimetic-S86,T172 mutant exhibited a synergistic effect, similar to the phosphomimetic-S41 mutant, whereas their phosphoablative counterparts did not (Figure 1D). Collectively, these findings reveal that the CaN-sensitive phosphosites S86 and T172 on GAP-43, identified through an unbiased phosphoproteomic screen in a model of α-syn pathology, play a key regulatory role in GAP-43 neurite branching.

### 2. GAP-43 phosphosites S86 and T172 regulate dendritic arborization and spine number in primary cortical neurons

We next asked if the effect on neurite branching, we observed in PC12 cells, can be recapitulated in primary cortical neurons. Rat cortical neurons were infected with lentiviruses expressing fusion proteins of GFP with either WT GAP-43, or the phosphomimetic and phosphoablative double mutants of S86,T172 phosphosites (hereafter referred to as phosphomimetic and phosphoablative double mutants). To visualize the dendritic arbor, we transfected neurons with the fluorescent protein mScarlet or stained them with BODIPY 558/568 Phalloidin. BODIPY 558/568 Phalloidin binds to F-actin, enabling fluorescence microscopy-based observation of actin fibers and, consequently, neuronal morphology. To assess neurite branching, we performed Sholl analysis. Expression of WT GAP-43 caused an increase in neurite branching compared to GFP alone (Figure 2A,B and Supplementary Figure 2A,B). The phosphomimetic double mutant further enhanced the dendritic arborization compared to WT GAP-43. Importantly, this effect was phosphorylation-dependent, as the phosphoablative double mutant did not show an increase in branching complexity relative to both, the WT and the phosphomimetic double mutant (Figure 2A,B and Supplementary Figure 2A,B).

**Figure 2.**
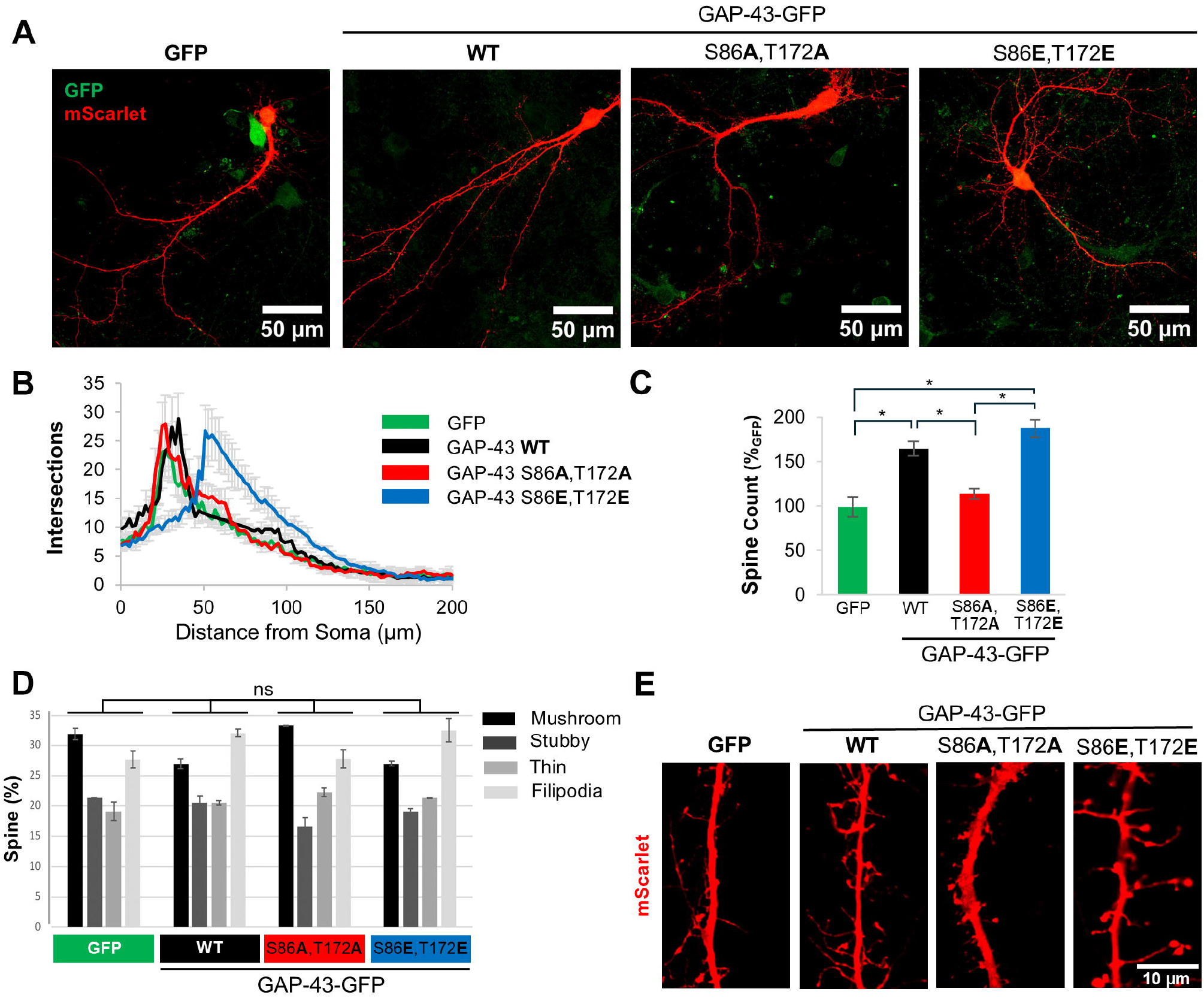
Calcineurin-dependent phosphosites S86 and T172 of GAP-43 contribute to neurite branching and spine formation in primary cortical neurons. **(A)** Representative confocal images of rat primary cortical neurons transduced with GFP or GFP fusions of GAP-43 WT, phosphoablative mutant (S86A,T172A), or phosphomimetic mutant (S86E,T172E) at the CaN-dependent phosphorylation sites. Cultures were transfected with a fluorescent protein mScarlet. Scale bar is 50µm. **(B)** Sholl analysis of the cultures in conditions presented in (A). N=30-50 cells. **(C)** Total number of spines/10µm from conditions in (A). **(D)** Distribution of spine morphology from conditions in (A). One-way Anova with post hoc Tukey’s test was used to analyze spine morphology distribution amongst the treatment groups. The distribution was not significantly different. **(E)** Representative images of dendritic spines from cultures in (A). Scale bar is 10µm. In all conditions, SEM and *p< 0.05 One-way Anova with post hoc Tukey’s test.

Given that the actin cytoskeleton was affected by GAP-43 CaN-dependent mutants, we investigated the effect of GAP-43 in dendritic spines, as they are highly dependent on actin dynamics. Expression of WT and phosphomimetic double mutant GAP-43 increased the total number of spines relative to phosphoablative double mutant and to GFP alone (Figure 2C). While the changes in spine number were dependent on S86,T172 phosphorylation, the morphological patterns (mushroom, stubby, filipodia and thin-like dendritic spines) were not significantly different between GFP, and WT and CaN-dependent phosphomutants of GAP-43 (Figure 2D,E). However, the distribution of GAP-43 within spines appears to be very different between the CaN-dependent phosphomutants (Supplementary Figure 2C). The phosphoablative double mutant exhibits lower expression and greater diffusion throughout the spine compared to the phosphomimetic double mutant and WT, both of which are highly expressed in the spines (Supplementary Figure 2C). Together, these data indicate that the CaN-sensitive sites S86,T172 are critical to promote branching and spine formation in primary cortical neurons.

### 3. GAP-43 phosphosites S86 and T172 mediate protein interactions enriched in ribosomal proteins

To identify protein interactors responsible for the neurite branching and spine effects of GAP-43 we took an immunoprecipitation (IP) MS-based approach, utilizing four different experimental conditions: WT, phosphoablative double mutant, phosphomimetic double mutant GAP-43-GFP and GFP alone to control for nonspecific protein copurification. HeLa cells were transfected with these constructs and IP using anti-GFP antibodies. GAP-43 IP were robust and, importantly, recovered similar amounts of GAP-43 across WT and the phosphomutants (Supplementary Table 1). Positive hits were considered those that either were present in the GAP-43 pulldowns and not in the GFP alone pulldowns or, if present in the GFP alone pulldown, those hits that were statistically significant to GAP-43-specific pulldowns. Under these criteria, we retrieved a total of 31 candidates for GAP-43-interacting partners that scored as positive hits. GAP-43 WT shared 6 protein interactors with the phosphoablative mutant, which were enriched for proteins involved in rRNA processing and structural components of ribosomes. In contrast, only one interactor, nucleophosmin (NPM1)—a protein involved in ribosome nuclear export—was shared between the WT and both mutants (Figure 3A,B). Additionally, 24 protein interactions were enriched in the phosphoablative mutant, which were also related to proteins involved in rRNA processing, structural ribosomal components, and cadherin binding (Figure 3A,B). Together, these findings suggest that the CaN-dependent phosphorylation sites mediate interactions with ribosomal proteins to regulate neurite outgrowth.

**Figure 3.**
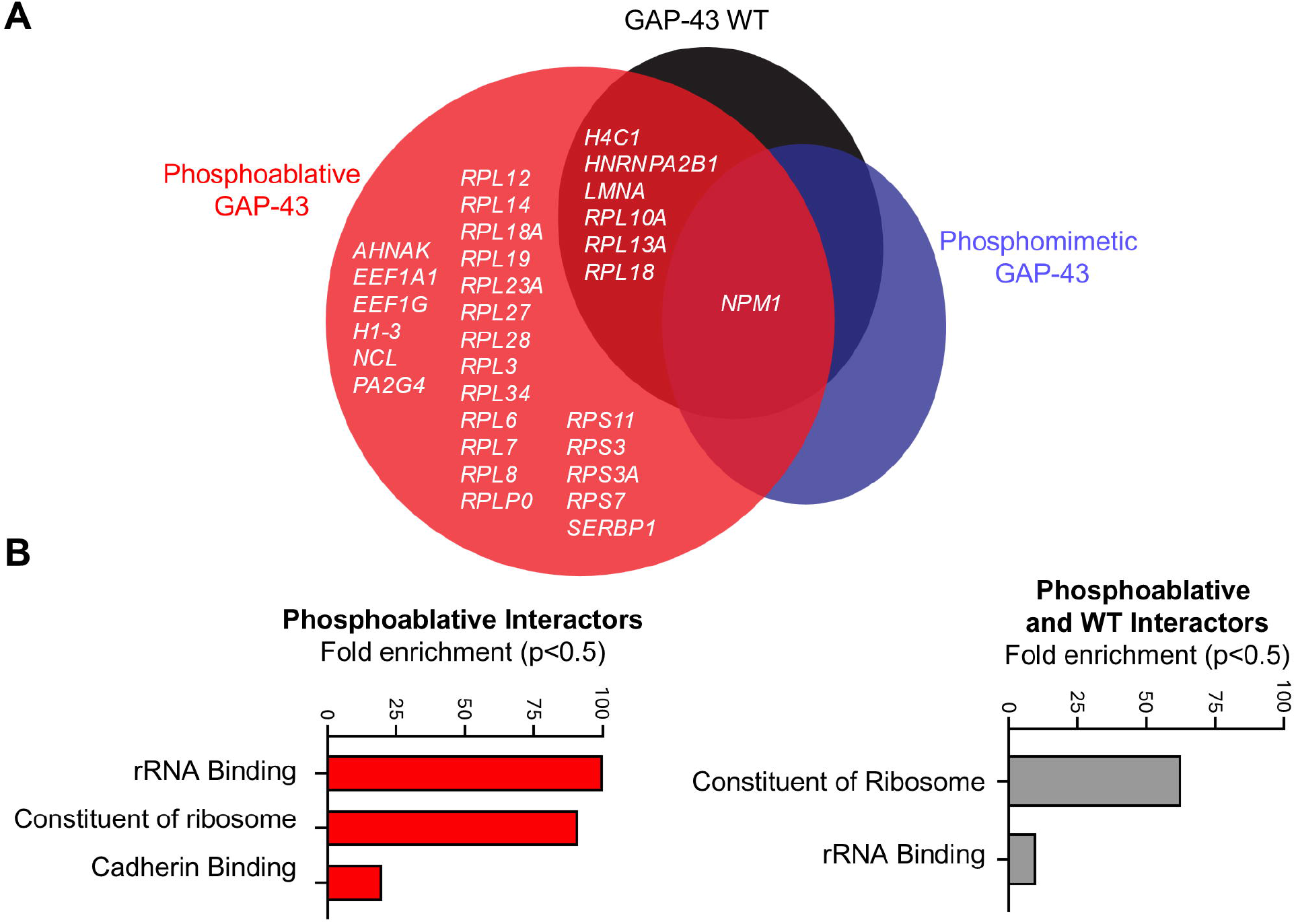
Calcineurin-dependent phosphosites S86 and T172 of GAP-43 bind structural components of ribosomes. **(A)** Venn diagram representation of all MS hits that passed the selection criteria described in main text and materials and methods. N = 3. One-Way Anova with a multi-comparison test using a two-stage linear step-up procedure of Benjamini, Krieger, and Yekutieli, q = 0.05. **(B)** Gene ontology analysis for molecular processes from the MS hits in (A).

### 4. The GAP-43 phosphomutants S86 and T172 can protect against α-synuclein neurodegeneration in primary cortical neurons

Phosphorylation at S86 and T172 was associated with neuroprotection in a large preclinical rat study of α-syn-expressing rats treated with sub-saturating doses of FK506 (Caraveo et al., 2017) (Figure 1A,B). To investigate if the effects of GAP-43 phosphomimetic double mutant in the dendritic arbor could protect against α-syn toxicity, we co-transduced primary cortical neurons with the disease-associated form of α-syn, A53T (Polymeropoulos et al., 1997), or with an empty vector as control, along with GAP-43 WT and CaN-dependent double phosphomutants (Figure 4A,B). Five weeks post infection, we immunostained the cultures for the neuronal specific marker MAP2 (microtubule-associated protein 2) to determine the number of neurons after each condition. In addition, we measured ATP levels as an alternative surrogate for viability. While overexpression of GAP-43 WT or GAP-43 phosphoablative double mutant did not have an effect in α-syn toxicity, as measured by MAP2+ neurons and ATP, overexpression of the phosphomimetic double mutant protected the neurons against the toxic effects of α-syn (Figure 4C-E). Moreover, the protective effects of the phosphomimetic double mutant were associated with an increase in PSD95 (postsynaptic density protein 95), a key scaffolding protein in excitatory synapses which levels serve as a surrogate marker for synaptic stability and maturation (Figure 4F,G). Together, these findings suggest that phosphorylation at the CaN-dependent sites S86 and T172 can influence α-syn pathology by regulating dendritic arborization and synapse stability.

**Figure 4.**
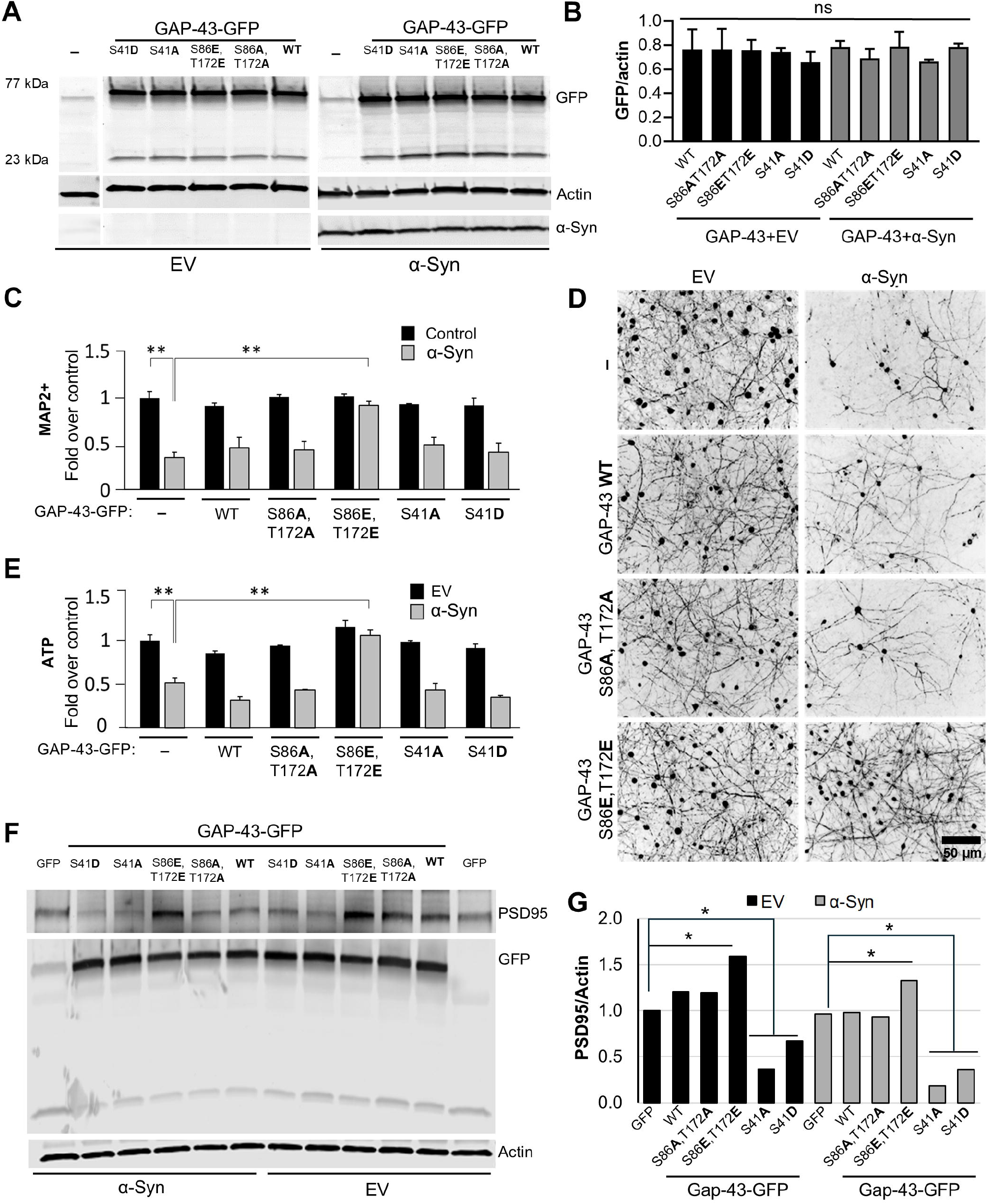
Calcineurin-dependent phosphosites S86 and T172 of GAP-43 regulate α-synuclein-mediated pathobiology. **(A)** Representative western blot from rat primary embryonic cortical neurons transduced with either GFP, or GFP fusions of GAP-43 Wild type (WT), phosphoablative (S86A,T172A) or phosphomimetic (S86E,T172E) mutants of the CaN-dependent phosphorylation sites, or the previously established PKC-dependent site (S41) in phosphoablative, S41A, or phosphomimetic, S41D, mutant form, in combination with Empty vector (EV) or αSyn A53T. Actin serves as a loading control. **(B)** Quantitation of GFP fluorescence intensity over actin from (A). SEM and *p< 0.05 One-way Anova with post hoc Tukey’s test. **(C)** Quantification of the neuronal marker MAP2 of the neuronal cultures immunostained for MAP2 **(D). (E)** Quantification of ATP levels from conditions in (A). Experiments in (A-E), N=3. **(F)** Representative western blot for PSD95, GFP and actin from cultures in (A) and quantitation of PSD95 signal/actin, the loading control **(G)**. One-way Anova with post hoc Tukey’s test; *<0.05; **<0.01; ***<0.001.

## Discussion

Using an unbiased phosphoproteomic approach and model of α-syn pathology in rats treated with FK506, we identified GAP-43 as one of the two primary substrates regulated by CaN in the striatum (Caraveo et al., 2017). Although these phosphorylation sites were previously identified in phosphoproteomic screens (Lundby et al., 2012) and linked to physiological neurite outgrowth (Biarc et al., 2012; Spencer et al., 1992), their causal role in regulating GAP-43 activity and α-syn neurodegeneration had not been explored. Here, we used a genetic phosphomutant approach to dissect the contribution of these phosphorylation events to neurite growth under physiological and α-syn conditions. Under physiological conditions, we demonstrated that phosphorylation of GAP-43 at S86 and T172 (mimicked by phosphomimetic mutants) increased neurite branching and spine density, whereas constitutive dephosphorylation by CaN (mimicked by phosphoablative mutants) reduced these features. Furthermore, we discovered that phosphorylation at these CaN-sensitive sites facilitates the recruitment of proteins involved in translation. More than half of the phosphoablative interactors are part of the large ribosomal subunit (RPL12, RPL14, RPL18, RPL19, RPL23A, RPL27, RPL28, RPL3, RPL34, RPL6, RPL7, RPL8, and RPLP0). A smaller fraction (∼16%) belongs to the small ribosomal subunit (RPS11, RPS3, RPS3A, and RPS7), while approximately 25% of the remaining interactors are involved in the regulation of mRNA translation (SERBP1, AHNAK, EEF1A1, EEF1G, NCL, and PA2G4) (Figure 3). Among these, Neuroblast differentiation-associated protein AHNAK (AHNAK), a Ca^2+^-regulated protein involved in cytoskeletal organization (Benaud et al., 2004; Shankar et al., 2010), and Elongation factor 1-alpha 1 (EEF1A1), a translation elongation factor whose activity is controlled by signaling and is also known to interact with actin and microtubules to regulate the cytoskeleton (Bosutti et al., 2022), are particularly interesting candidates for further investigation. Previous studies have established that the ribosomal apparatus is essential for protein synthesis, which supports dendritic growth and maintenance (Slomnicki et al., 2016; Jaworski et al., 2005; Kwon et al., 2011; Lesiak et al., 2013; Kumar et al., 2005; De Rubeis et al., 2013). Particularly, GAP-43 and actin mRNAs were found to be locally translated in regenerating axons (Kalinski et al., 2015). Additionally, the downregulation of ribosome signaling and eukaryotic translation initiation factor 2 pathway genes following GAP-43 overexpression suggests a role for GAP-43 in regulating cellular translation (Chen et al., 2021). Based on our current interactome data, we propose that phosphorylation at S86 and T172 is a signaling mechanism that enables recruitment of the translation machinery and proteins that activate dormant mRNAs, facilitating local protein synthesis at growth cones to promote dendritic and spine growth. Further studies will be needed to fully elucidate the role of GAP-43 in mRNA regulation at growth cones.

We previously showed that dephosphorylation of GAP-43 at S86 and T172 was associated with α-syn-induced neurodegeneration, whereas phosphorylation at these sites correlated with neuroprotection in a large preclinical rat model of α-syn pathology treated with FK506 (Caraveo et al., 2017) and (Figure 1A,B). Here, using rat primary cortical neurons we established a causal relationship for this association. Under α-syn pathological conditions, phosphorylation of GAP-43 at S86 and T172 (mimicked by phosphomimetic mutants) conferred neuroprotection against α-syn proteotoxic stress whereas the phosphoablative mutant did not (Figure 4C-E).

Regulation of GAP-43 has been explored as a therapeutic strategy for neurodegenerative diseases. Specifically, the effects of agents that upregulate GAP-43, such as BDNF (brain-derived neurotrophic factor), have been tested in animal models of PD (Gupta et al., 2009). In one PD patient, infusion with glial cell-derived neurotrophic factor at the putamen provided benefits even after cessation of treatment (Patel et al., 2013; Palasz et al., 2020). Treatments with Pilose antler extracts in a rat model of PD increased striatal GAP-43 protein expression and reduced dopaminergic SNc neuronal cell death (Li et al., 2019). Despite the positive effects of neurotrophic factors in animal models, their short half-life, low bioavailability, and limited permeability through the blood-brain barrier (BBB) imposed challenges in their application to patients (Palasz et al., 2020). Challenges associated with using neurotrophic factors to elevate GAP-43 levels can be circumvented by employing pharmacological agents capable of crossing the BBB and modulating GAP-43 activity, such as through phosphorylation. A promising candidate is the FDA-approved CaN inhibitor FK506.

The neuroprotective effects of FK506 on neurite growth in both PD and non-PD models have been well documented for over two decades (Butcher et al., 1997; Gold, 1997; Snyder et al., 1998; Madsen et al.,1998; Zuber and Donnerer, 2002). FK506 exerts its effects by inhibiting CaN through the formation of a tertiary complex with the immunophilin FKBP12. Studies have demonstrated that FKBP12 inhibition alone significantly contributes to neuroprotection in PD models (Steiner et al., 1997; Costantini et al., 1998; Tanaka & Ogawa, 2004; Gerard et al., 2010) and plays a role in α-syn pathobiology by preventing its aggregation (Gerard et al., 2006; Gerard et al., 2008). Other studies have also highlighted the role of CaN inhibition in neuroprotection (Zaichick and Caraveo, 2023; Martin et al., 2012; Mondal et al., 2023). Our previous findings addressed this controversy by demonstrating that CaN interacts with FKBP12, enabling the dephosphorylation of substrates (Caraveo et al., 2017). In the context of α-syn pathobiology, we showed that constitutive activation of the CaN/FKBP12 complex leads to the dephosphorylation of substrates, such as GAP-43, which in the presence of α-syn contributes to neurodegeneration. However, partial inhibition of CaN/FKBP12 with FK506 alters CaN’s substrate specificity (due to differences in enzyme-substrate affinity), which in the context of α-syn results in neuroprotection. Additionally, we demonstrated that FKBP12 inhibition, independent of CaN, also confers neuroprotection against α-syn toxicity (Caraveo et al., 2017). These findings explain why cyclosporin A, another CaN inhibitor, is not as effective against

α-syn proteotoxic stress. The elucidation of FK506’s dose-dependent explains why the maximal neuroprotection against α-syn can only be achieved with sub-saturating doses of FK506, when both CaN-dependent and CaN-independent pathways are engaged (Caraveo et al., 2017). While FK506 is commonly administered at saturating doses in clinical settings to prevent organ rejection—where full CaN inhibition is necessary—our proposed sub-saturating doses, which induce only partial CaN inhibition, would likely result in minimal and transient immunosuppression, given the drug’s circulating half-life of approximately 12 hours (Venkataramanan et al., 1995). At sub-immunosuppressive doses (10-fold lower than standard clinical doses), the risk of adverse effects such as opportunistic infections, posterior reversible leukoencephalopathy, and seizures—commonly associated with FK506 at immunosuppressive levels—would be significantly reduced. Notably, epidemiological studies have reported a lower prevalence of Parkinson’s disease in patients prescribed FK506 (Silva et al., 2024), further supporting its potential therapeutic role.

This study demonstrated that one of FK506’s neuroprotective effects is exerted through the modulation of GAP-43 phosphorylation, thereby promoting neurite outgrowth and mitigating α-syn proteotoxicity. Sub-saturating FK506 doses would finely tune GAP-43 to drive sufficient, but not hyperactive, dendritic sprouting in a spatially and temporally confined manner. This regulation is necessary to promote regeneration specifically in the affected neuronal populations at the right time window while preventing potential complications arising from hyperconnectivity. Further exploration of these areas using human-derived neuronal lines will be crucial to corroborate the effects of GAP-43 S86 and T172 phosphomimetic mutants and facilitate the development of GAP-43-targeting therapies for neurodegenerative diseases.

## Materials and Methods

### 1 Plasmids and viruses

Lentiviral Plasmids: pLV-αSyn A53T (hSynapsin promoter) was a generous gift from Aftabul Haque and Susan Lindquist (Whitehead Institute for Biomedical Research, Cambridge, MA). Control (empty vector) plasmid was generated by excising αSyn and ligating the resulted product. Murine GAP-43-FLAG pAG plasmid was a generous gift from Eric Norstrom (DePaul University Chicago). The GAP-43 mutagenesis was performed using Q5 Site-Directed Mutagenesis Kit (E0554, New England Biolabs) according to the manufacturer protocol. Murine GAP-43-FLAG pcDNA 3.1/Hygro construct was used as a template DNA. T95, S86 and T172 were substituted with either A or EE; S41 was substituted with either A or DD.

For the experiments with primary cortical neurons, GAP-43 WT and single phosphoablative and phosphomimetic mutants S41A and S41DD (referred to as S41D in the main text), as well as double phosphomutants S86EE,T172EE (referred to as S86E,T172E in the main text) and S86A,T172A were subcloned into the lentiviral expression vector pLV-eGFP. Viruses were produced in the laboratory and in the NU Gene editing transduction and Nanotechnology Core. In the laboratory, packaging plasmids (pMD2.G and psPAX2) were obtained from Addgene. Lentiviral constructs were packaged via lipid-mediated transient transfection of the expression constructs and packaging plasmids (pMD2.G and psPAX2) into HEK293T cells using a modified protocol from Park et al. (Park et al., 2008). Lentiviruses were purified and concentrated using the LentiX Concentrator (Clontech, PT4421-2) according to the manufacturer’s protocol. Lentivirus titer was determined using the QuickTiter Lentivirus titer kit (lentivirus-associated HIV p24; Cell Biolabs, VPK-107) or HIV-1 p24 ELISA pair set (Sino Biological Inc, SEK11695) according to the manufacturer’s protocol. All titers were further normalized in SH-SY5Y cells infected with gradual multiplicities of infection (MOI) volumes of lentivirus and processed for Western blot analysis 3 dpt. Equal MOIs were used to infect rat cortical cultures on 5DIV. SH-SY5Y cells are used for technical validation rather than biological experimentation. Specifically, they serve to test the expression of the lentiviruses used to transduce primary cortical neurons. This approach allows us to titrate the multiplicity of infection (MOI) to ensure that the constructs of interest are expressed at appropriate levels. This is particularly important in cases where the MOI between different lentiviruses does not align with the expression levels of the target protein—in this case, wild-type (WT) and phosphomutant variants of GAP-43.

### 2 Cell lines

PC12 cells were maintained up to passage 10 in DMEM high-glucose media supplemented with 10% NuSerum and 1% FBS (fetal bovine serum). 70-80% confluent culture was passaged by trypsinization (0.05% Trypsin/EDTA) followed by centrifugation at 500×g for 5 min and resuspension in full culture medium. For the evaluation of the response to GAP-43 overexpression, the cells were seeded on collagen-coated tissue culture polystyrene. 1 ml aqueous solution containing 35 µg/ml human placenta collagen IV (Advanced Biomatrix) and 15 µg/ml chicken collagen II (Sigma) was added to each well of a 24-well plate followed by overnight air-drying at 25 °C and UV-sterilization. Lipofectamine 3000-assisted transfection was performed 24 h prior to examination. For a single well of 24-well plate, 500 ng pcDNA 3.1/Hygro construct (EV, GAP-43-FLAG WT, phosphoablative S86A,T172A mutant, or phosphomimetic S86EE,T172EE mutant) and 83 ng pEGFP-C1 were added to 50 µl Opti-MEM along with 0.8 ml P3000 reagent. 0.8 µl of Lipofectamine 3000 reagent was diluted in 50 µl Opti-MEM in a separate sterile Eppendorf tube. Ater 5 min incubation, the DNA and Lipofectamine 3000 reagent solutions were combined, incubated for 30 min and then added to the PC-12 cells precultured for 24 h. After 5 h, the solution was replaced with the full growth media. Cells were imaged using Leica DMI3000B microscope fitted with an QImaging QIClick CCD Camera and Leica HCX PL FLUOTAR L 20×/0.40 CORR PH1 objective. Image acquisition was done using Q-capture pro7 software and manual tracking of the equal exposure and digital gain setting between images. All image processing and analysis was done using Fiji. All experiments were done four independent times with n>300 cells per condition.

HeLa and HEK293T (human embryonic kidney cells) cells were cultured at 37°C and 5% CO_2_ in Dulbecco’s Modified Eagle Medium (DMEM) with 4.5% glucose (Corning), 10% FBS (Denville) and 1× penicillin and streptomycin (Gibco).

### 3 Primary Cortical Cultures

Embryonic rat cortical neurons were isolated from euthanized pregnant Sprague–Dawley rats at embryonic day 18 using a modified protocol from Lesuisse and Martin (Lesuisse and Martin, 2002, Pacifici and Peruzzi, 2012). Protocol was approved by Northwestern University administrative panel on laboratory animal care. Embryos were harvested by Cesarean section and cerebral cortices were isolated and dissociated by 0.25% Trypsin without EDTA (Invitrogen, 15090-046) digestion for 15 min at 37°C and trituration with 1 ml plastic tip. Poly-D-Lysine (Sigma, P-1149)-coated 96-well and 24-well plates were seeded with 4 × 10^4^ and 2 × 10^5^ cells correspondingly in neurobasal medium (Invitrogen, 21103-049) supplemented with 10% heat-inactivated FBS (Gibco), 0.5 mM glutamine (Gibco), penicillin (100 IU/mL), and streptomycin (100 μg/mL) (Gibco). Before seeding, cells were counted using the Automated cell counter TC10 (Bio-Rad) and viability (90-95%) was checked with Trypan Blue Stain (0.4%, Gibco 15250-061). After 1 h incubation at 37°C, media was changed to neurobasal medium (Invitrogen, 21103-049) supplemented with B27 (Invitrogen 17504044), 0.5 mM glutamine, penicillin (100 IU/mL), and streptomycin (100 μg/mL). One-half (out of 100 µl volume for 96-well plates and 500 µl volume for 24-well plates) of the media was changed 4 h before transduction on day 5 *in vitro* (5DIV). As a surrogate marker of cell viability, cellular ATP content was measured two-three times a week between 5-7.5 weeks post transduction using the ViaLight Plus kit (Lonza, LT07-221) or ATPlite kit (PerkinElmer, 6016941) according to the manufacturer’s instructions.

### 4 Western blot

Infected neuronal cultures (5w2d post transduction) were lysed using a radioimmunoprecipitation (RIPA) assay buffer (50mM Tris/HCl pH 7.6; 150mM NaCl; 20mM KCl; 1.5mM MgCl_2_; 1% NP40; 0.1% SDS). In all experiments, lysis buffer was supplemented with the Halt protease and phosphatase inhibitor cocktail (Thermofisher; 78441). Samples were incubated on ice for 30 minutes and pushed through a 27G needle (10 times) to ensure full lysis. Samples were then centrifuged at 21,000×g for 20 minutes and the obtained supernatants were used for Western blot analysis. Protein concentration was analyzed with the Pierce BCA Protein Assay kit (Thermofisher) and Fisherbrand™ accuSkan™ GO UV/Vis Microplate Spectrophotometer (Fisher Scientific). After the addition of the appropriate amount of the 6× Laemmli Sample Buffer (Bioland scientific LLC, sab03-02) with 5% ß-mercaptoethanol (Sigma) protein samples (10-30 µg) were boiled and separated on precasted 4-20% Criterion TGX Stain-free gels (Bio-Rad) and transferred to a nitrocellulose membrane (Amersham Protran 0.2um NC, #10600001). Membranes were blocked with 5% non-fat milk in Tris-buffered saline (TBS) (50 mM Tris, pH 7.4, 150 mM NaCl) for 1 h at 25°C. Membranes were subsequently immunoblotted overnight in primary antibodies (anti-α-syn, BD, 610787; anti-Actin, Abcam, ab6276; anti-PSD95, Thermofisher, MA1-046) at 4°C, shaking. The following day, the membranes were washed three times with TBST (TBS with 0.1% Tween) for 5 minutes each and incubated in secondary IRDye antibodies for 1 h shaking at 25°C. Membranes were washed three times with TBST before imaging using Li-Cor Odyssey® CLx Imaging System. Images were processed using Image Studio Software (LI-COR Biosciences) and signal densities were quantified using Fiji (Schindelin et al., 2012).

### 5 Immunofluorescence

Transduced neuronal cultures (4-5 weeks post transduction, 96-well plates) were fixed with 3% (vol/vol) paraformaldehyde. Cells were then washed three times with PBS for 5 minutes each wash and permeabilized using in 0.1% Triton X-100 in PBS (PBST), at 25°C for 20min and then blocked in 2% Bovine Serum Albumin (BSA, Sigma, SLBT8252) in PBST for at least 60 min at RT. Primary antibodies (anti-MAP2, Millipore, AB5622) were diluted in the blocking buffer and incubated overnight at 4°C followed by triple wash and incubation (1-2 h at 25°C) with secondary antibodies, also diluted in the blocking buffer. Subsequently, cells were washed trice and incubated with Hoechst (Invitrogen, 33342) for 10 min at 25°C followed by triple rinsing with PBS. Cells were imaged using Leica DMI3000B microscope fitted with an QImaging QIClick CCD Camera and Leica HCX PL FLUOTAR L 20×/0.40 CORR PH1 objective. Image acquisition was done using Q-capture pro7 software and manual tracking of the equal exposure and digital gain setting between images. All experiments were done three independent times. Images were recorded from at least 4 wells and analyzed in a randomized and blinded fashion. MAP2-positive cell bodies (neurons) were manually calculated and subtracted from the total nuclei count (Hoechst). Fluorescence quantification was done using a Microplate Reader Infinite® M1000 PRO (Tecan) using appropriate excitation and emission wavelengths. Images were processed using ImageJ/FIJI software built-in package for color deconvolution (ABC-DAB) and processed for average intensity. Intensity values over six fields for each sample are compared and processed for statistical analysis.

### 6 Scholl analysis and spine determination

Scholl analysis was performed in 21DIV primary rat cortical neurons. The cells were visualized by BODIPY 558/568 Phalloidin staining and transfection with the fluorescent protein mScarlet. Prior to phalloidin staining, the neurons were fixed with 4% paraformaldehyde in PBS for 15 minutes at 25 °C. Subsequently, the cells were washed with PBS trice for 5 min each and permeabilized with 0.1% Triton X-100 in PBS for another 15 minutes. Then the cells were washed twice with PBS and stained for one hour at 25 °C in the dark with 165nM BODIPY 558/568 Phalloidin prepared in PBS with 1% BSA. After the staining, the cells were washed with PBS trice for 5 min each, immunostained for GFP, and mounted on coverslips. The neurons were imaged using Nikon A1R GaAsP point-scanning laser confocal microscope using 63× oil-immersion objective (NA = 1.4) with z-series of 8-10 images, taken at 0.3 μm intervals, with 1024×1024 pixel resolution. The stacks were then flattened to 1 image in ImageJ and Scholl Analysis performed using the SNT plugin (ImageJ) after manual tracing. Morphometric analysis was performed on spines from two dendrites (secondary or tertiary branches), totaling 100 μm, from each neuron. Single dendritic spines were determined from the intensity of BODIPY 558/568 Phalloidin. Mushroom-shaped spines were considered as head/neck ratio >1.1 and spine diameter >1 micron (spine area>0.78µm^2^). For each condition, n=30-50 neurons.

Lipid-mediated transient transfection was performed using Lipofectamine 2000 (Thermofisher, L3000015). 1.2 µg pmScarlet3_C1 (Addgene, 189753) as well as 3.7 µl of Lipofectamine 2000 were added to 120 µl fresh neurobasal media in separate sterile Eppendorf tubes, then incubated for 5 min and combined for another 20 min at 25 °C and added to a 35 mm FluoroDish (World Precision Instrument FD35100) with cultured neurons transduced with either GFP or GAP-43-GFP viruses. 3 hours post-transfection, the media was replaced with conditioned media for another 24 h followed by fixation with 4% paraformaldehyde and 4% sucrose in PBS for 15 minutes at 25 °C.

### 7 Immunoprecipitation (IP)

HeLa cells were transiently transfected with pER4-GFP and GAP-43 (WT and double phosphomutants S86A,T172A and S86D,T172E) subcloned in a pER4-GFP vector. Lipid-mediated transient transfection was performed using Lipofectamine 2000 (Thermofisher, L3000015) according to the manufacturer’s protocol. 24 hours post-transfection, the cells were briefly washed with ice-cold PBS and lysed using 1% Triton X-100, 10% glycerol, 10 mM Tris/Cl pH 7.5; 150 mM NaCl, 0.5 mM EDTA. Lysis buffer was supplemented with the Halt protease and phosphatase inhibitor cocktail (Thermofisher, 78441). Samples were incubated on ice for 30 minutes and pushed through a 27G needle (10 times) to ensure full lysis and then centrifuged at 21,000×g for 20 minutes. Protein concentration was analyzed with the Pierce BCA Protein Assay kit (Thermofisher). Each supernatant was incubated rotating at 4°C for 2 h with 30 ul of 50% slurry GFP-Trap beads (ChromoTek GFP-Trap® Agarose beads) prepared in lysis buffer. Next, the beads were washed trice in a centrifuge (2,500×g, 2 min, 4°C) with lysis buffer and two more times – with the wash buffer (10mM Tris-Cl pH7.5, 150mM NaCl, 0.5mM EDTA). IP samples were submitted for TMT-MS at the MIT Kock Proteomics facility.

### 8 Statistical Analysis

One-way ANOVA with Tukey’s post hoc test was used for three or more dataset quantifications. Statistical calculations were performed with GraphPad Prism 7 Software (http://www.graphpad.com), p values <0.05 were considered significant. To correct for multiple comparisons in MS data analysis, the two-stage linear step-up procedure of Benjamini, Krieger and Yekutieli post-hoc tests was used to control the false discovery rate (FDR) and assess differences between phosphomutants, q = 0.05. Results are expressed as average + the standard error mean (SEM). Regression analysis was done using Microsoft Excel Statistics add-in package. A minimum of 3 independent biological replicates were used for each experiment with at least 3 replicates per sample within each experiment. The specific number of biological replicates for each experiment is provided in the figure legends.

