## Supplementary Figure 1 for "Calcineurin-mediated regulation of growth-associated protein 43 is essential for neurite and synapse formation and protects against α-synuclein-induced degeneration"

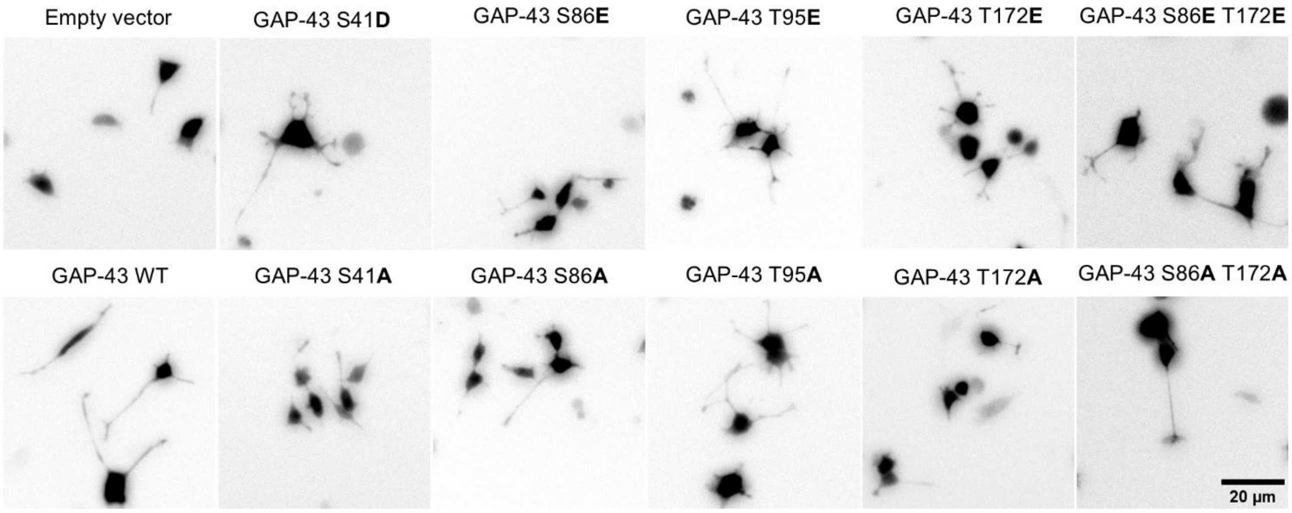


**Supplementary Figure 1. Calcineurin-dependent phosphosites S86 and T172 of GAP-43 contribute to neurite branching in PC12 cells.** Representative confocal images of PC12 cells co-transfected with GFP for neurite visualization and GAP-43 either WT, phosphomimetic mutants S41D (as a positive control), S86E, T95E, T172E or S86E-T172E double mutant, and phosphoablative mutants S41A, S86A, T95A, T172A or S86A-T172A double mutant. 20×; scale bar is 20µm.
