## Supplementary Figure 2 for "Calcineurin-mediated regulation of growth-associated protein 43 is essential for neurite and synapse formation and protects against α-synuclein-induced degeneration"

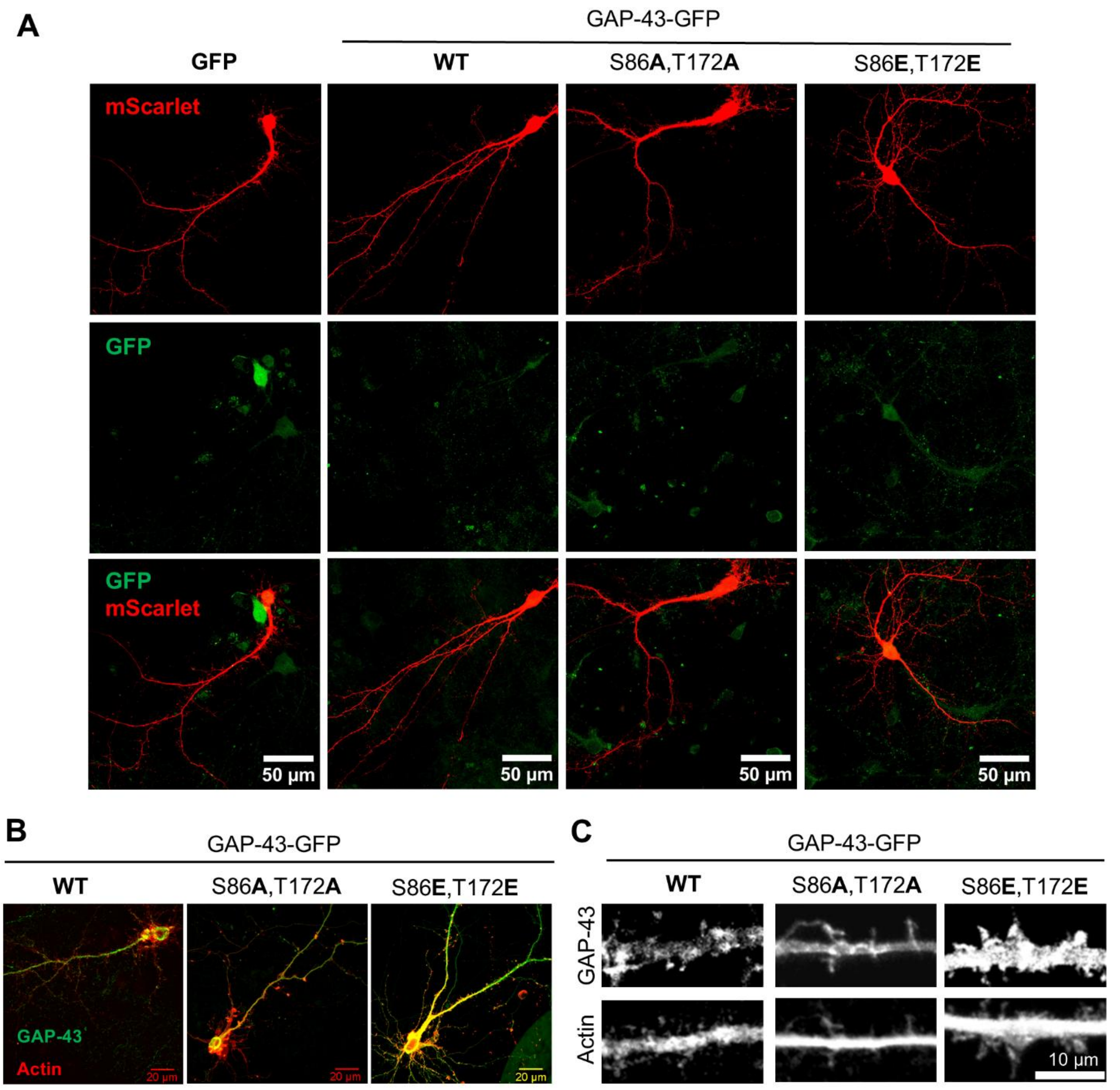


**Supplementary Figure 2. Calcineurin-dependent phosphosites S86 and T172 of GAP-43 contribute to neurite branching and spine formation in primary cortical neurons.**
**(A)** Representative confocal images of primary rat cortical neurons transduced with GFP or GFP fusions of GAP-43 WT, phosphoablative double mutant (S86A,T172A), or phosphomimetic double mutant (S86E,T172E) at the CaN-dependent phosphorylation sites and transfected with mScarlet to visualize neuronal morphology. Scale bar is 50µm. **(B)** Representative confocal images of rat primary cortical neurons transduced with GFP or GFP fusions of GAP-43 WT, phosphoablative mutant (S86A,T172A), or phosphomimetic mutant (S86E,T172E) at the CaN-dependent phosphorylation sites. Cultures were immunostained for GFP (green channel) and BODIPY 558/568 Phalloidin-stained for actin (red). Scale bar is 20µm. **(C)** Representative images of dendritic spines from cultures in (B). Scale bar is 10µm.
